# The effects of chronic social stress on cognitive flexibility in adult female macaques

**DOI:** 10.1101/2025.08.01.667938

**Authors:** MC Alvarado, T Jonesteller, K Bailey, AC Gray, MM Sanchez, J Bachevalier

**Affiliations:** Emory National Primate Research Center, Emory University, Atlanta, GA; Department of Psychiatry & Behavioral Sciences, Emory Univ., Sch. of Med., Atlanta, GA; Department of Psychology, Emory University, Atlanta, GA

**Author notes:** **CORRESPONDING AUTHOR:** Maria Alvarado, PhD, Emory National Primate Research Center, Emory University, 954 Gatewood Rd., NE, Atlanta, GA 30329.

**Keywords:** Intradimensional/Extradimensional task, temperament traits

## Abstract

Chronic social subordination stress in macaques, particularly beginning early in life, is associated with negative health and cognitive aging outcomes. Utilizing a longitudinal, translational monkey model of early life stress in Rhesus macaques (*Macaca mulatta)*, we assessed adult female cognitive performance that had received social subordination stress associated with Low Social Status (LSS) since birth. This chronic social stress in lower ranked monkeys produces physiological stress responses that over time can accelerate biological and cognitive aging. We compared 14 adult females of Low Birth Rank that had received higher levels of social subordination stress since infancy with 11 adult females of High Birth Rank that had received lower levels of chronic social stress. We followed these two groups longitudinally from birth to adolescence to assess long-term behavioral, physiological, and neural consequences of social stress. As they reached adulthood (ages 7-8 years), we measured executive function/cognitive flexibility using the Intra-/Extra-dimensional shift task (ID/ED), relevant for age-related cognitive decline. Only mild differences between High and Low ranking subjects were observed on the simple discrimination, and the reversal learning stages of the task. We did find an interaction between High Birth Rank and performance across the three dimensional-shift stages that was not present in the low-ranking subjects. Thus, although at this first age, we did not yet detect the presence of accelerated cognitive decline following chronic social stress, the low-ranking females showed mild deficits in cognitive flexibility compared to high-ranking subjects. We discuss additional factors impacting performance, and comparisons with neuroimaging data.

## 1. INTRODUCTION

Chronic social stress, especially beginning early in life and continuing through the lifespan, drives neuroinflammatory states and evokes epigenetic changes that negatively affect brain developmental trajectories (Cohen et al., 2012; Lupien et al., 2018). This chronic stress results in structural and functional alterations of brain structures, such as the prefrontal cortex (PFC), hippocampus (HIPP) and amygdala (AMY), at different developmental time points leading to accelerated cognitive decline (Fjell and Walhovd, 2010; Morrison and Baxter, 2012; Simen et al., 2011). Animal models with short life spans (invertebrates, rodents) have provided valuable information on the impact of early stress on accelerated brain and cognitive changes at later age, but they have significant limitations to inform about human aging and the therapeutics developed from those models have failed in clinical trials. Nonhuman primates (NHP) by contrast provide a translational model of cognitive aging due to their longlife span, their wide range of cognitive abilities, as well as gradual aging-related cognitive impairments and brain pathology similar to those in humans (Didier et al., 2016).

Age-related cognitive decline in NHPs is characterized by deficits in executive functions, cognitive flexibility and memory that emerged gradually, starting in middle-aged animals and deteriorating further at older ages. In a cross-sectional study of normally aging NHPs (3 to 31 years of age), age-related impairments for visuospatial abilities emerged between 16-21 years, yet recognition memory impairments were not detected until the late 20’s (Bachevalier et al., 1991; Presty et al., 1987). Additional NHP studies have indicated that one of the first executive functions to be impacted in middle-aged subjects was behavioral and cognitive flexibility, usually measured with reversal or attentional shift tasks (Bonté et al., 2011; Gullstrand et al., 2022; Herndon et al., 1997; Lacreuse et al., 2018; Lai et al., 1995; Moore et al., 2005; Voytko, 1999). Interestingly, high individual variability was also observed for the older groups in almost all previous studies, suggesting that some animals exhibited accelerated cognitive decline, whereas others performed as well as younger animals. These findings have not only been reported in rhesus macaques (see for review, Baxter et al., 2023; Rapp and Amaral, ;1992; Zeamer et al., 2011), but also in several species of NHPs, such as baboons (Lizarraga et al., 2020), African green vervet monkeys (Varma et al., 2024) and marmosets (Sadoun et al., 2019; Vanderlip et al., 2024; Vanderlip et al., 2023). The individual variability of cognitive performance in the aged animals is interesting and indicates that the impact of intrinsic factors (such as genetics) but also of extrinsic factors (such as environmental or social stress) can affect the developmental trajectory of the brain and of the aging cognitive decline. Finally, almost all the earlier observations on aged-NHPs came from cross-sectional studies but only few prospective longitudinal studies have followed the animals from adulthood to aging (e.g. (Rothwell et al., 2022), and none have investigated longitudinally the effects of early social stress on brain aging and cognitive decline.

We recently initiated a prospective longitudinal aging study based on our previous developmental work (NIH-funded studies (R01 HD077623) using Rhesus macaques to assess the lifelong effects of chronic social subordination stress early in life on physiological markers, brain neuropathology and, socioemotional and cognitive changes associated with accelerated aging. We used a naturally occurring social stressor displayed by macaques to maintain social hierarchies, social subordination. Social subordination in female macaques is a well-validated model for chronic psychosocial stress (Howell et al., 2014; Wilson, 2016) that produces stress-related phenotypes in adults (Jarrell et al., 2008; Kaplan et al., 2010; Michopoulos et al., 2012a, 2012b; Shively, 1998; Shively et al., 1997; Snyder-Mackler et al., 2016; Tung et al., 2012). Because Rhesus macaque infants inherit the relative social rank of their mothers (Bernstein, 1976), we studied newborn infants from birth to menarche that were born into the social rank of their dam and fell into one of two groups: infants from Low-Social Status (Submissive) mothers (SUB: Low-SS) and infants of High-Social Status (Dominant) mothers (DOM: High-SS). We found that SUB: Low-SS subjects showed several early physiological markers of cellular aging, accelerated DNA methylation age, shortened telomere and inflammation, detectable from infancy to early adulthood (Drury et al., 2017). We have also reported neural changes that included enlarged AMY and HIPP volumes (Kovacs-Balint et al., 2018; Kovacs-Balint et al., 2025; Kovacs-Balint et al., 2024a, 2024b; Kyle et al., 2019; Pincus et al., 2021), reduced NAA in anterior cingulate cortex, and presence of amyloid-beta levels (Alzheimer’s disease markers (Lesuis et al., 2018)) that emerged at the juvenile and early adolescent ages. In the present study, we continued investigation of this unique cohort of macaques with early chronic social subordination stress and characterized their cognitive functions as they have reached full adulthood (7-8 years). Our preliminary brain data have already shown that, as they reached adulthood (7-11 years), the larger AMY volume observed in the SUB-Low-SS females between 6 months and adolescence, was still evident in adulthood. Also, as compared to the DOM-High-SS adult females, the SUB-Low-SS adult females had larger orbitofrontal cortex, medial PFC and HIPP volumes as well as greater negative connectivity between PFC area 9 and the AMY and the HIPP (Kovacs-Balint et al., 2024b; Kovacs-Balint et al., 2021).

Given the brain regions impacted by chronic social stress, particularly experienced early in life, the present study tested these now SUB-Low SS and DOM-High SS adult females on the Intra- /Extra-dimensional Attentional Shift Task (ID/ED), measuring behavioral and cognitive flexibility known to be affected by aging processes (Amelchenko et al., 2023; Bonté et al., 2011; Gullstrand et al., 2022; Kupis et al., 2021; Lacreuse et al., 2018; Moore et al., 2003; Moore et al., 2005; Mota et al., 2019) but see also Zeamer et al., 2011) that will serve as a baseline rom which future testing may reveal differing rank-based aging trajectories of cognitive decline.

However, given their changes in brain structure and connectivity, we anticipated that some Low-ranking females from birth might already show signs of impaired attentional flexibility as compared High-SS females. In addition, we speculated that the presence of chronic social stress could be at least one factor that may explain the high individual cognitive scores reported in the aged populations in almost all studies of cognitive decline in NHPs (Abbott et al., 2003). But we also assessed other factors that may have impacted the performance of the animals in the ID/ED task. These factors included the animals’ temperament traits when interacting in their social environment and after training on the task, the group size in which the animals were living, or the diet they had ingested as infants and juveniles.

## 2. METHODS

### 2.1 Subjects, Experimental Groups and Housing

Subjects were 25 adult female rhesus macaques (*Macaca mulatta)* aged 7-8 years at the start of this study and naïve to cognitive testing. They were born and raised in large, multi-family social groups at the Emory National Primate Research Center (ENPRC) Field Station breeding colony in Lawrenceville, GA, an AAALAC Accredited facility. Currently, they live in social groups of varying size, from 8-180 adults and their offspring. Groups are generally composed of several multigenerational matrilines, with 1-4 adult breeder males. Animals are provided ad lib water, daily chow (Purina Mills Int., Lab Diets, St. Louis, MO, USA) supplemented with a variety of fresh vegetables, daily oranges as well as forage and destructible enrichment. Animals have access to outdoor and indoor spaces. As part of the breeding colony, they continued to have yearly infants if the group had a breeder male and they successfully gave birth. When new births occurred, those subjects were given a month at least away from testing to bond with their newborn. Testing then resumed if the subject was cooperative and they brought their infant with them during testing until they were old enough to stay behind (6 months old or more). Otherwise, infants of these subjects were not part of the study. The protocols for this study were approved by the Emory University Institutional Animal Care and Use Committee (IACUC) in accordance with the Animal Welfare Act and the U.S. Department of Health and Human Services “Guide for Care and Use of Laboratory Animals”. These animals were assigned in two testing cohorts, with half starting the first year (Cohort 1) and the other half the second year (Cohort 2) of the study. Each cohort had almost equal numbers of each social ranking Table 1 summarizes the demographic of all animals, including age, diet they received from infancy through puberty, social rank at Birth and at the time of this study.

**Table 1.**

| Subject<br>Birth ID* | Cohort | Prepubertal<br>Diet | Social Rank |  |  |  | Rank<br>Changes | Adult<br>Group<br>Size | Test<br>Age<br>(Years) |
| --- | --- | --- | --- | --- | --- | --- | --- | --- | --- |
|  |  |  | Birth Rank |  | Cog Rank |  |  |  |  |
|  |  |  | Percentile | H/L | Percentile | H/L |  |  |  |
| DOM-1 | 1 | HCD | 86.05 | High | 92.78 | High | 0 | 180 | 8.42 |
| DOM-2 | 1 | LCD | 72.09 | High | 67.01 | High | 0 | 180 | 8.25 |
| DOM-3 | 1 | LCD | 71.95 | High | 68.75 | High | 0 | 45 | 8.25 |
| DOM-4 | 1 | LCD | 69.77 | High | 62.89 | High | 0 | 180 | 8.17 |
| DOM-5 | 1 | HCD | 62.67 | High | 50.00 | High | 2 | 25 | 7.42 |
| DOM-6 | 1 | HCD | 67.07 | High | 50.00 | Low | 2 | 45 | 7.42 |
| DOM-7 | 1 | LCD | 60.98 | High | 18.75 | Low | 1 | 45 | 7.33 |
| DOM-8 | 1 | HCD | 71.95 | High | 81.25 | High | 0 | 45 | 7.42 |
| DOM-9 | 2 | LCD | 72.00 | High | 70.00 | High | 0 | 30 | 8.08 |
| DOM-10 | 2 | LCD | 80.23 | High | 73.20 | High | 0 | 180 | 6.83 |
| DOM-11 | 2 | HCD | 76.74 | High | 82.47 | High | 0 | 180 | 6.75 |
| SUB-1 | 1 | HCD | 18.60 | Low | 25.00 | Low | 2 | 8 | 8.42 |
| SUB-2 | 1 | HCD | 18.60 | Low | 25.00 | Low | 2 | 15 | 8.42 |
| SUB-3 | 1 | HCD | 5.81 | Low | 28.57 | Low | 3 | 12 | 8.33 |
| SUB-4 | 1 | LCD | 22.09 | Low | 75.00 | High | 1 | 12 | 8.33 |
| SUB-5 | 1 | LCD | 19.51 | Low | 0.00 | Low | 2 | 8 | 8.33 |
| SUB-6 | 1 | HCD | 24.42 | Low | 75.00 | High | 2 | 8 | 8.33 |
| SUB-7 | 2 | LCD | 24.00 | Low | 24.00 | Low | 1 | 8 | 7.92 |
| SUB-8 | 2 | HCD | 37.33 | Low | 0.00 | Low | 2 | 8 | 7.75 |
| SUB-9 | 2 | LCD | 20.00 | Low | 14.29 | Low | 0 | 15 | 8.00 |
| SUB-10 | 2 | HCD | 1.22 | Low | 36.36 | Low | 1 | 45 | 7.92 |
| SUB-11 | 2 | LCD | 20.00 | Low | 0.00 | Low | 2 | 8 | 7.08 |
| SUB-12 | 2 | LCD | 3.66 | Low | 0.00 | Low | 0 | 30 | 7.17 |
| SUB-13 | 2 | HCD | 10.67 | Low | 0.00 | Low | 0 | 15 | 6.92 |
| SUB-14 | 2 | HCD | 17.33 | Low | 50.00 | High | 1 | 15 | 6.83 |

Note: Demographic information for the 25 rhesus macaque females studied and divided in two cohorts aged 7-8 years at the time of cognitive testing. The Subject ID reflects the original Birth Ranks (DOM or SUB). From birth to end of puberty, half of the subjects in each cohort, had been fed a typical, low calory diet (LCD), whereas the other half had access to both the LCD diet and to a highly caloric diet (HCD). Social ranks: the table provides the rank each animal had been assigned to at birth and then at the time of cognitive testing. At each time point, social ranking was determined from observations of subordinate/submissive behaviors in the group (Low, and High) and by calculating the status of their family relative to all families in their group (which is presented as a percentile), such that percentiles closer to zero represent lower social status and percentiles closer to 100 represent higher status. Finally, because several animals have changed social rank since birth due to changes in the composition and sizes of their social groups, we also provided the number of times each animal has changed social ranking. Additional details in text.

### 2.2 Previous studies on these subjects

As mentioned above, in infancy, these animals contributed to a larger developmental study (birth to puberty) that examined the effects of early life social subordination stress and obesogenic diet on brain structural and functional development (neuroimaging), emotional reactivity, stress neuroendocrinology (hypothalamic-pituitary-adrenal –HPA-axis regulation) and inflammation (see published data (Korobkova et al., 2023; Kovacs-Balint et al., 2025; Kovacs-Balint et al., 2024a, 2024b; McCormack et al., 2022; Morin et al., 2022; Morin et al., 2020; Morin et al., 2019; Sanchez et al., 2015). Each animal was born either to a high-ranking, or dominant (DOM) matriline, or to a low-ranking, or subordinate (SUB) matriline. From birth through puberty within each social ranking group, half were fed a typical, low calorie diet (LCD; Purina Mills Int., Lab Diets, St. Louis, MO, USA), whereas the other half had postnatal access to both the LCD diet and to a highly caloric diet (HCD; (4.47 kcal/g, D07091204S Research Diets, New Brunswick NJ)) (See Table 1 for prepubertal Diet assignments). After puberty and the conclusion of that study, all animals were reassigned to the ENPRC breeding colony care, remaining in their natal social groups, and from that time, all have received the standard LCD diet.

### 2.3 Social Rank

Fourteen of these females were Low ranking at birth (Subordinate (SUB) group, 6 LCD and 8 HCD), and 11 of them were high ranking at birth (Dominant (DOM) group, 6 LCD and 5 HCD). Ranking information was provided by ENPRC Colony Management staff based on observations of dyadic agonistic interactions, first at birth by using the social ranking of their dams and at the time of this study when the animals reached adulthood, and also routinely verified by our own group observations 3 times annually. We also routinely conferred with ENPRC Colony Management for observations or reports from their staff suggesting individuals were trying to assume higher ranks. Following established methods (Bernstein, 1976; Embree et al., 2013; Howell et al., 2014), an adult female was considered as ranking below a groupmate if they invariably responded submissively when that individual approached and/or acted aggressively towards them. Social rank was thus determined from observations of subordinate/submissive behaviors, such as withdrawing, grimacing, and fleeing, rather than aggressive behaviors. Rhesus infants inherit their mother’s matrilineal social rank/status, which reflects the status of their family/matriline relative to all families in the group/home compound. For example, a rank of 2/8 would indicate the infant was born into the second-ranking family out of eight total families in that group and is thus a relatively high-ranking individual. Birth Ranks were normalized to a percentile by dividing the family rank by the total number of families, then subtracting this value from one (e.g., 2/8 = 75%), such that percentiles closer to zero represent lower social status and percentiles closer to one represent higher status.

#### 2.3.1 Lifetime Rank changes

A main goal of this study was to determine the long-term consequences of chronic social stress that began early in life, which in this model reflects the cumulative experienced social stressors due to social Rank. However, prior to and during the present study, a number of adult females had a shift in social rank, due to changes in the dynamic of the social group when group tensions required the movement of some matrilines to smaller social groups (see Table 1 for details). As we were interested, for this report, in the impact of low social status on successful cognitive testing, we compared whether subjects’ Birth Rank played the more important role in adult cognitive performance than adult rank at the time of testing (Cog Rank to differentiate between lifetime and current effects of Rank. For use as a continuous variable, this was calculated as the individual’s percentile rank (using the same dyadic interaction scale) with respect to the total group, rather than family (e.g., 15^th^/180, or 8^th^/9 individuals) and normalized as described above. For use as a grouping variable, we utilized “High” or “Low” to designate subjects with percentile ranks >50, with the exception of subjects that held ranks of 50% in small groups but still outranked other group members at the same percentile, in which case the dominant one of the two was ranked “High” (see Table 1).

### 2.4 Cognitive testing

Monkeys were trained on the Intra-/Extra-dimensional (ID/ED) discrimination task, which is an analog of the Wisconsin card sort task (Dias et al., 1996), measuring cognitive flexibility, mediated by the prefrontal cortex (PFC), parietal cortex and striatum (Berry et al., 2016; Braver et al., 2003; Klanker et al., 2013; Kupis et al., 2021). It assays two types of cognitive control: (i) attention towards aspects of the environment that are relevant for reinforcement by requiring subjects to shift attention away from previously relevant stimulus dimension (e.g. shape) towards a newly relevant dimension (e.g. line), that is an extra-dimensional shift (EDS) or a shift of attention from previously relevant stimuli of the same dimension (shape, such as a square and circle) towards newly relevant stimuli of the same dimension (triangle and rectangle), that is an intra-dimensional shift (IDS) and (ii) the capacity to learn that a previously reinforced stimulus is now not-reinforced (reversal shift). Several studies have shown that the EDS depends on the integrity of the dorsolateral PFC and the reversal shift depends on the orbital frontal cortex (Burnham et al., 2010; Dias et al., 1996; Fernandez-Ruiz et al., 2001; Rogers et al., 2000) whereas the IDS depends on the integrity of the striatum (Berry et al., 2016; Fernandez-Ruiz et al., 2001; Klanker et al., 2013). After pre-training sessions, the task was presented in several stages of increasing difficulty (see below for details).

#### 2.4.1 Apparatus

Testing was conducted in a Wisconsin General Testing Apparatus (WGTA), located inside a darkened, sound-shielded room. Additional sound masking was provided by a white-noise generator. The stationary test tray (30 cm X 63 cm) contained three, 25 mm diameter food wells that were spaced 5 cm apart and aligned 13 cm from the front of the animal’s cage. Vertical screens were raised and lowered to hide the testing tray from the monkey during intertrial intervals, and to hide the experimenter from the monkey when making its choice. These subjects are group housed, active breeders, therefore food restriction was never used. Rather, rewards for correct choices included ENPRC approved treats such as flavored pellets, cubed fruits or vegetables, raisins, etc. Food preferences per subject were noted on testing logs to ensure use of the most appealing rewards for each subject.

#### 2.4.2 Testing Scheduling

Testing occurred 3-4 days per week, depending upon subject rank, social group dynamics and comfort with the testing environment, breeding and birthing seasons, as well as other project-related data collections not reported here.

#### 2.4.3 Habituation and Pre-Training

Before beginning the ID-ED task, all monkeys were first re-trained to be accessed from their social groups (a familiar procedure at the ENPRC), following procedures in place for over 40 years to minimize arousal (Wilson et al., 2013), and adverse health effects (Blank et al., 1983; Walker et al., 1982; Wilson et al., 1986). They were then transported to the testing suite to be acclimated to the room, the personnel, testing cage and the apparatus and then were trained to displace objects covering the food wells to retrieve food rewards. This acclimation/habituation consisted of five phases. In *Phase 1*, monkeys were transported from their housing group to a cage in the testing suite with which they were familiar. The cage contained a bowl filled with food rewards and an experimenter was in proximity. When monkeys retrieve the rewards from the bowl without hesitation, they moved to *Phase 2*, where they were acclimated to taking treats from the trainer until they consistently did so without hesitation. In *Phase 3*, they were placed into the testing cage in the testing room and gradually moved into the WGTA, with the 2 screens raised and allowed to retrieve rewards scattered on the tray until they retrieved the rewards without signs of fearfulness. During this shaping, the opaque screen separating the monkeys from the tray and the one-way mirror screen separating the tray from the experimenter were also lowered progressively few inches at a time from trial to trial until monkeys were comfortable with the movement of the screens and retrieve rewards on each trial with the human screen closed. In *Phase 4,* an object was introduced on the tray positioned first close to the food well and then covering the well progressively until monkeys retrieved food rewards from wells entirely covered. Finally, in *Phase 5,* the animals were pretrained on a daily simple-discrimination using two 3-D objects. When animals displaced the correct objects on 5-6 trials in a row, they were moved to the formal testing on the ID/ED task.

#### 2.4.4 ID/ED task

White plastic plaques with a clear top served as stimulus picture holders (10 cm wide X 12.5 cm long) into which different printed pictures of colored shapes could be inserted (9.5 cm wide X 10 cm long). The task was presented in several stages (Fig. 1) formed by a series of simultaneous visual discrimination problems in which a response to one of two stimuli resulted in a food reward. For each stage, animals received a maximum of 30 trials per daily session, separated by 15-sec intervals. *Stage 1*: Simple Discrimination (SD): response to one of the two aqua colored shapes was reinforced until criterion was met. *Stages 2-4*: are a series of three simple reversals (SR) of the reward contingency learned in Stage 1. That is, on the first reversal (SR1), the monkey needed to select the stimulus that was *not r*ewarded in Stage 1. In the second and third reversals the reward will be again switch from one stimulus to the other. *Stage 5*: Compound Discrimination (CD) included the aqua shapes onto which orange lines were superimposed, creating compound stimuli with Shape and Line dimensions, but only Shape predicted reward. *Stage 6*: Intra-Dimensional Shift (IDS): a new pair of compound stimuli (new Shapes, new Lines) were presented, with one of the two new Shapes predicting reward (i.e. shift within the dimension of Shape). *Stage 7*: Extra-Dimensional Shift (EDS): Novel shapes and lines stimuli were again introduced, but now the line dimension was relevant and were reinforced while the shape dimension was redundant and irrelevant. It should be noted that in primate studies the ID/ED task usually includes reversal discrimination problems after each stage to test for behavioral inhibition; however, to maintain the task procedures as similar as possible to those used for testing humans for closer species comparisons, in the present study we opted to include three reversal discrimination problems only after the simple discrimination of Stage 1. . Criterion for each Stage was set at 10 correct responses in a row and data recorded include the number of total trials and errors to criterion for each stage. Animals were allowed a maximum of 900 trials to achieve criterion. If they failed at any stage, they did not continue to the remaining stages.

**Figure 1:**
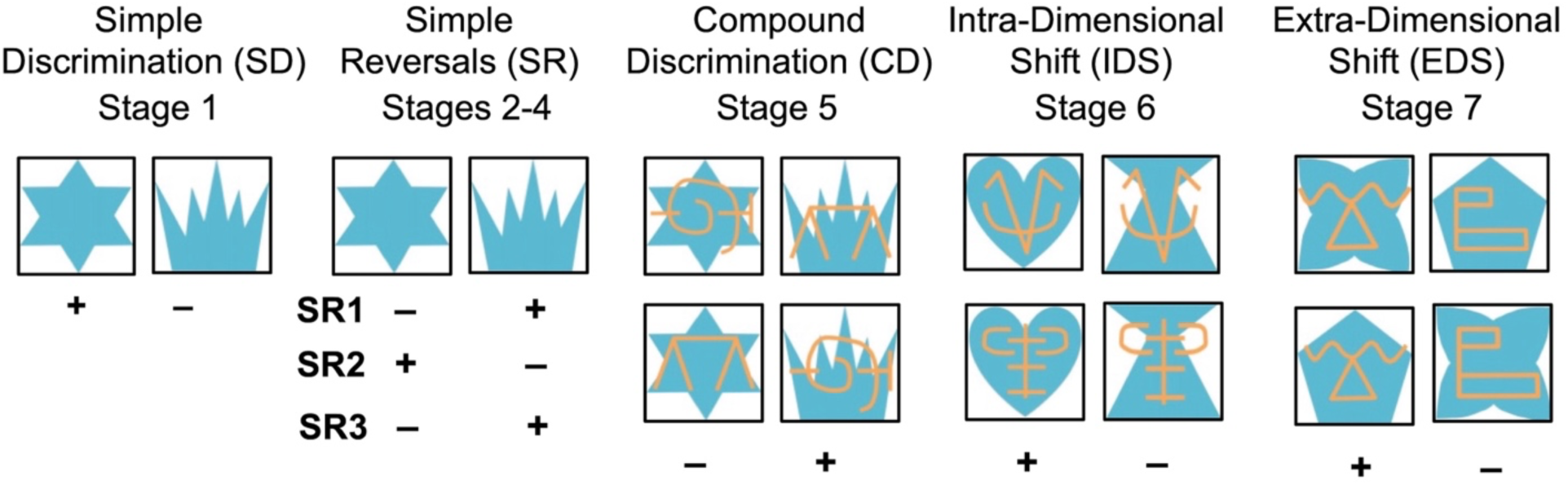
Schematic of Intra-dimensional/Extra-dimensional Shift task (ID/ED). Depicted are the stimulus pairs for each stage. The Left-Right position of each correct stimulus was equally reinforced and determined pseudorandomly. The +/− beneath stimuli indicate the rewarded stimulus for each pair. For Stages 5-7, there are two discrimination problems, depicted by the upper and lower pairs. Only one pair was presented on each trial, but the presentation order was predetermined and intermixed pseudorandomly. See text for further details. Adapted from (Weiss et al., 2019).

Likewise, if a subject refused to participate further at any point, and despite remedial efforts (e.g., giving them some time off, changing treats etc.) stopped choosing stimuli, their testing was discontinued, and their scores were only counted for fully completed Stages. *Correction procedure*: If at any point an animal developed a strong side-bias, responding only to one side for at least 5 testing days, a correction procedure was introduced during which training continued, but on each trial the rewarded stimuli was always placed on the side opposite to the animal’s spatial bias (e.g. left food well for an animal with right bias). When the animal changed its strategy and displaced the rewarded stimulus for 5 trials in a row, regular training was resumed. These correction trials were not included in the data presented.

### 2.5 Personality Trait Assessments

To better understand how/whether temperamental variables can impact successful cooperative testing of socially housed macaques, we also conducted personality assessments of each monkey, both within their social groups, averaged across multiple observations (“Ob-traits”) and a single assessment by the primary cognitive tester encompassing all completed stages for each monkey (“Cog-traits”). The assessment was derived from the Hominoid Personality Questionnaire assessments published for large social groups (Weiss et al., 2011) and adapted for our subjects. For this paper, we utilized personality traits that could be scored both in the testing environment and the social group. This resulted in 19 behaviors that were compared across the two settings. Of these 19, only 12 were very likely to yield scores in both settings and were selected. Table 2 lists those 12 personality traits with a brief definition for each. For each monkey, each behavior was subjectively rated by one of three experimenters with extensive expertise in primate social and emotional behavior, using a 7-point Likert-type scale. The definitions used for each Likert-scale level were as follows: 1-Displays either total absence or negligible amounts of the trait; 2 - Displays small amounts of the trait on infrequent occasions., 3 - Displays somewhat less than average amounts of the trait., 4 - Displays about average amounts of the trait, 5 - Displays somewhat greater than average amounts of the trait, 6 - Displays considerable amounts of the trait on frequent occasions, 7 - Displays extremely large amounts of the trait. Scores between observers/testers were discussed and evaluated amongst the three observers, to ensure consistency in applying scores in both “Ob” and “Cog” settings.

**Table 2:** Summary of Temperament Composite Scores (adapted from Weiss et al, 2011)

| *Traits* | *Definitions* |
| --- | --- |
| *Fearful* | • Subject reacts excessively to real or imagined threats by displaying behaviors such as screaming, grimacing, running away or other signs of anxiety or distress. |
| *Cautious* | • Subject often seems attentive to possible harm or danger from its actions, avoids risky behaviors. |
| *Timid* | • Subject lacks self-confidence, is easily alarmed and is hesitant to venture into new situations. |
| *Anxious* | • Subject often seems distressed, troubled, or is in a state of uncertainty. |
| *Gentle* | • Subject responds to tester in an easy-going, kind, and considerate manner; not rough or threatening. |
| *Cool* | • Subject seems unaffected by emotions, is usually undisturbed, assured, and calm |
| *Curious* | • Subject has a desire to see or know about objects, devices, or other monkeys. |
| *Aggressive* | • Subjects often initiates fights or other menacing and agonistic encounters with other monkeys or the tester. |
| *Irritable* | • Subject often seems in a bad mood or is impatient and easily provoked to anger exasperation and consequent agonistic behavior. |
| *Defiant* | • Subject is assertive or contentious; maintains these actions despite unfavorable consequences. |
| *Excitable* | • Subject is easily aroused to an emotional state. Becomes highly aroused by situations that would cause less arousal in most monkeys. |
| *Erratic* | • Subject is inconsistent, indefinite, and widely varying in its behavior and moods. |

Note: Personality traits adapted from Hominoid Personality Questionnaire after Weiss et al., (Weiss et al., 2011). This subset was utilized as raters were able to score these traits in both the cognitive testing (Cog) and social observation (Ob) settings. Traits were scored using a Likert Scale with the following rating definitions: 1. Either total absence or negligible amounts of the trait; 2. Small amounts of the trait on infrequent occasions; 3. Somewhat less than average amounts of the trait; 4. About average amounts of the trait; 5. Somewhat greater than average amounts of the trait; 6. Considerable amounts of the trait on frequent occasions; 7. Extremely large amounts of the trait. The row borders indicate traits that tended to correlate with each other in the cognitive testing ratings (See also Table 4)

## 3. DATA ANALYSES

### 3.1 Rank Grouping Variable

As shown in Table 1, many subjects in this study have experienced changes in social rank such that their Birth Rank differs from their current Cog Rank. Although correlations between rank at birth and rank at cognitive testing revealed them to be highly correlated using either High/Low rankings (r_s_(23) = .60, p = .002) or Percentile (r_s_(23) = .68, p < .001), we nevertheless found for some measures, the impact of Birth Rank (and therefore High/Low Early Life Social Stress) were better predictors of task performance, whereas Cog Rank was a better predictor of temperament traits during habituation to testing. Because Birth Rank may represent the impact of programming factors during the developmental period that may impact later behavioral and cognitive functions, whereas Cog Rank may represent cumulative stress factors at the time of testing, we therefore, present separate results using Birth and Cog Rank as the main grouping factors for all data analyses.

### 3.2 Habituation/Pretraining and ID/ED performance

The number of Phases completed for the habituation and pretraining were summed and compared by Birth and Cog Rank, using a non-parametric Mann-Whitney U test. Trials and Errors to criterion were recorded for all 7 Stages, but to avoid ceiling effects, only Errors were compared for each ID/ED Stage. In addition, given that the stages measure different cognitive abilities, we divided the stages into 3 components and compared between Rank (Birth and Cog). Thus, scores on Stage 1, which measured abilities to learn stimulus-reward associations, was analyzed separately using one-way ANOVA with Rank as factor. Two-way repeated measure ANOVAs (RM-ANOVA) including Birth and Cog Ranks as the between subject factor and stages as repeated measures within subject factor were used to analyze the effects of rank on the three Reversal stages (Stages 2-4), which measured shift in reward contingency, as well as on the CD, IDS, and EDS stages (stages 5-7), which required a shift in relevant stimulus contingency. Finally, we also calculated the number of perseverative errors each animal made during each of the 3 reversals and used a two-way repeated measure ANOVAs (RM-ANOVA) including Birth and Cog Ranks as the between subject factor and reversals as repeated measures within subject factor to measure behavioral inhibition.

### 3.3 Personality Traits

During habituation and testing, we observed differences in certain animals’ willingness to participate in behavioral testing. In particular, High-ranking animals (Cog Rank) were less willing to engage with testers even for their favorite food rewards. By contrast, Lower-ranking subjects were more readily engaged by food rewards, possibly due to the lack of social competition in the testing setting. In addition, we had observed personality differences among High- and Low-ranking animals from our observations in their social groups. Therefore, we also conducted personality assessments of each subject at the completion of cognitive testing and compared the impact of Birth or Cog Rank (as a categorical or percentile variables) and Personality Traits that could predict the likelihood of a subject successfully participating in cognitive testing. To avoid using multiple highly correlated factors, we first conducted Spearman Rank correlations to select individual, non-correlating measures, or to create composite factors for highly correlated measures (see Supplementary Figures 1-3). These composite scores (which were averages of the scores across the traits in each grouping) were used as factors in the Exploratory Associations (see below in Results section).

#### 3.3.1 Correlations by context

We first compared associations between traits scored in either context (Cog-traits or Ob-traits) with themselves (Supplementary Figures 1-2) and then with each other (Supplementary Figure 3). For those Cog-traits, we found strong positive correlations among the Fearful_Cog_, Anxious_Cog_, Cautious_Cog_ and Timid_Cog_ traits, which we used to create a composite score called “Timid_Cog_” by averaging across the four traits for each subject. A second cluster can be seen with strong positive correlations between the Aggressive_Cog_, Irritable_Cog_, Defiant_Cog_, Excitable_Cog_ and Erratic_Cog_ traits, which we used to form a “Bold_Cog_” composite score. A third set included Gentle_Cog_, Cool_Cog_ and Curious_Cog_ that we used to create a composite score called “Calm_Cog_”. In addition to these positive correlations, strong negative relationships were observed between the Curious_Cog_ trait and each of the traits comprising the Timid cluster, confirming the observation that Anxious or Fearful subjects were likely negatively impacted by the novelty of objects or the testing environment and therefore less curious. Similarly, the Aggressive_Cog_ and Irritable_Cog_ traits correlated negatively with those in the Timid cluster, (except for Anxious), and the Gentle trait correlated negatively with the Aggressive_Cog_, Irritable_Cog_, Defiant_Cog_ and Excitable_Cog_ traits. All Spearman’s rho and p values reported in Supplementary Figure 1.

For Ob-traits, scored in social housing (see Supplementary Figure 2), we found similar, but not identical relationship among the traits. Positive correlations were observed among the traits used to create the “Timid_Cog_” cluster for the Cog setting. We created a “Timid_Ob_” composite score for those traits. Similarly, moderate positive correlations were found among the Aggressive, Irritable, Defiant and Erratic (unlike Cog, it did not include the “Excitable” trait), which we combined to form a “Bold_Ob_” cluster. The traits comprising the “Calm_Cog_” cluster were not correlated in the group observations and so were included as single traits. As was observed for the Cog traits, the Aggressive_Ob_ trait was moderately to strongly negatively correlated with the components of the “Timid_Ob_” traits, as well as with the “Gentle_Ob_” trait. Lastly, the “Cool_Ob_” trait was negatively correlated with the “Anxious_Ob_” trait. All Spearman’s rho and p values reported in Supplementary Figure 2.

#### 3.3.2 Correlations between context

As shown in Supplementary Figure 3, comparisons between composite Cog-trait scores (Timid_Cog_, Bold_Cog_, and Calm_Cog_) and composite Ob-trait scores (Timid_Ob_, Bold_Ob_) as well as the Gentle_Ob_, Cool_Ob_, and Curious_Ob_ and Excitable_Ob_ non-clustered traits showed a strong positive correlation for Timid across the two conditions and a moderate positive correlation between Gentle_Ob_ (single trait) and Calm_Cog_ composite score. All Spearman’s rho and p values are reported in Supplementary Figure 3.

### 3.4 Exploratory association analyses

In preliminary analyses of the data, we explored the contribution of several additional variables to determine their impact on the willingness of animals to engage or succeed at cognitive testing. Predictive measures used as covariates in ANOVA analyses described above included Prepubertal Diet, given that approximately half of the subjects were reared with the opportunity to consume a dense calorie diet for their first two years of life (Table 1) and that this dietary difference between animals has resulted in volume changes in brain areas important for cognition (Kovacs-Balint et al., 2025; Kovacs-Balint, Jonesteller, et al., 2024; Kovacs-Balint, Wang, et al., 2024). In addition, we correlated (Spearman Rank) measures such as *Group Size* with performance on the various phases of Cognitive testing (e.g. Habituation, Priming, Errors etc. during Simple Discrimination, the Reversals or the Shifts). However, as none of these measures influenced the data obtained with the ANOVA analyses or correlated highly with Rank, they were not reported in the Results section below.

## 4. RESULTS

### 4.1 Habituation and Pretraining

Performance during the Pretraining phase is shown in Table 3. Subjects are listed by the social ranks measured at Birth (Birth Rank), and also included is each subjects’ current rank at the time of testing (Cog Rank). Ranks are indicated by group (High/Low) or as their percentile rank. The number of Sessions to complete Habituation, and Trials to pass Priming are listed, with group averages calculated by Birth and Cog Ranks. Of the 25 subjects trained, 5 did not complete the habituation or pretraining for varied reasons. Two High Birth Ranked subjects gave birth during the habituation Phases and quit participating (of these two, one was High and one Low Cog Rank), 3 others (2 High, 1 Low (Birth Ranks same as Cog Ranks)) were either not motivated by treats, or never acclimated to the environment, and so did not progress to formal testing. *Analyses by Birth Rank*. High ranking subjects on average required 22.89 Habituation Sessions and, 38.14 Priming Trials. Low Ranking subjects on average required 25.93 Sessions, and 23.23 Trials. Mann-Whitney U comparisons revealed no difference by Rank on performance [U=54 and 30.5, respectively; all p’s>0.05]. *Analysis by Cog rank*. High Cog ranking subjects required 22.17 Sessions, and 34.60 Trials on average, whereas Low-ranking subjects required 27.55 Sessions, 22.30 and Trials. No effect of Cog Rank was observed on these measures [U= 47.0, and 33.0 respectively, all p’s>0.05].

**Table 3:** Pretraining Performance.

| *Subject*<br>*Birth ID* | *Birth Rank* |  | *Cog Rank* |  | *Pretraining* |  |
| --- | --- | --- | --- | --- | --- | --- |
|  | *Percentile* | *Group* | *Percentile* | *Group* | *Habituation*<br>*Sessions* | *Trials to pass*<br>*Priming* |
| *DOM-1* | 86.05 | High | 92.78 | High | 20 | 32 |
| *DOM-2* | 72.09 | High | 67.01 | High | 17 | 26 |
| *DOM-3* | 71.95 | High | 68.75 | High | 19 | 24 |
| *DOM-4* | 69.77 | High | 62.89 | High | 24 | 128 |
| *DOM-5* | 62.67 | High | 50.00 | High | 27 | – |
| *DOM-6* | 67.07 | High | 50.00 | Low | – | – |
| *DOM-7* | 60.98 | High | 18.75 | Low | 36 | 24 |
| *DOM-8* | 71.95 | High | 81.25 | High | – | – |
| *DOM-9* | 72.00 | High | 70.00 | High | 14 | 19 |
| *DOM-10* | 80.23 | High | 73.20 | High | 33 | – |
| *DOM-11* | 76.74 | High | 82.47 | High | 16 | 14 |
| *SUB-1* | 18.60 | Low | 25.00 | Low | 21 | 10 |
| *SUB-2* | 18.60 | Low | 25.00 | Low | 16 | 36 |
| *SUB-3* | 5.81 | Low | 28.57 | Low | 13 | 26 |
| *SUB-4* | 22.09 | Low | 75.00 | High | 13 | 42 |
| *SUB-5* | 19.51 | Low | 0.00 | Low | 16 | 23 |
| *SUB-6* | 24.42 | Low | 75.00 | High | 21 | 24 |
| *SUB-7* | 24.00 | Low | 24.00 | Low | 29 | 12 |
| *SUB-8* | 37.33 | Low | 0.00 | Low | 42 | 14 |
| *SUB-9* | 20.00 | Low | 14.29 | Low | 26 | 11 |
| *SUB-10* | 1.22 | Low | 36.36 | Low | 36 | 13 |
| *SUB-11* | 20.00 | Low | 0.00 | Low | 35 | 19 |
| *SUB-12* | 3.66 | Low | 0.00 | Low | 23 | – |
| *SUB-13* | 10.67 | Low | 0.00 | Low | 45 | 54 |
| *SUB-14* | 1.22 | Low | 50.00 | High | 27 | 18 |
|  |  |  |  | ***Birth High*** | **22.89** | **38.14** |
|  |  |  |  | ***Birth Low*** | **25.93** | **23.23** |
|  |  |  |  | ***Means:*** |  |  |
|  |  |  |  | ***Cog High*** | **22.17** | **34.60** |
|  |  |  |  | ***Cog Low*** | **27.55** | **22.30** |

Note: Pretraining Performance: Number of Sessions to pass Habituation (Phases 1-4), Number of Trials to pass Pretraining (Phase 5). Em dashes indicate failure to advance to Testing (e.g., 4 High- and 1 Low-ranking animal failed to advance to formal testing). The Subject ID reflects the original Birth Ranks (Dom or Sub). Four subjects held a different rank at adult test from infancy (one lost ranking, 3 gained rank). Birth and Current Cog Ranks are listed by Percentile and High/Low. *Means* indicates the averages across Birth Rank and Cog Rank. See text for details.

### 4.2 ID/ED Testing

Performance on each phase of ID/ED testing is shown in Figure 2. Because we were interested both in the long-term consequences of social rank stress as well as in the impact of current rank on performance, the data are organized by Birth Rank (Fig 2A) and by Cog Rank (Fig 2B)..

**Figure 2:**
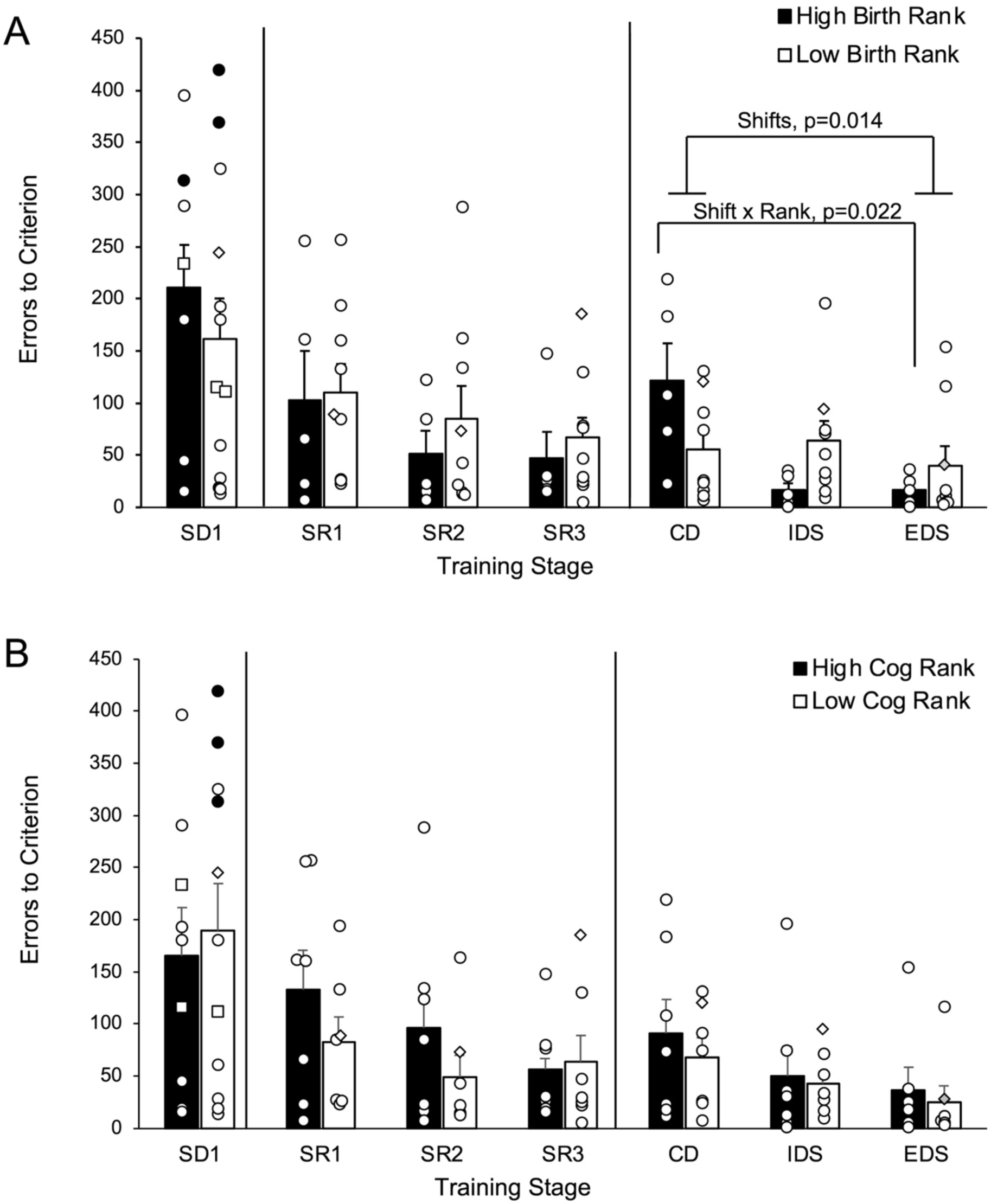
Errors to Criterion across the seven testing Stages x Social Rank. Panel A plots the data according to Birth Rank, Panel B plots the data according to Cog-Rank. Conventions for both panels: Individual data are shown (open circles). Data for subjects failing to reach criterion in 900 trials are shown in filled circles. Three subjects quit testing during the reversals; their data only contributed to SD1 (shown in open squares). Finally, one subject fell ill after IDS and was removed from the study. Those data are indicated in the open diamond and the data at Stage 7 was computed as the Group Mean for that Rank (grayed in diamond). Error bars equal SEM. Asterisks indicate p<0.05. See Results for explanation.

#### 4.2.1 Simple Discrimination (SD1)

*By Birth Rank*: As shown in Figure 2A, one High-ranking subject and two-Low ranking subjects failed to reach criterion, and their errors on this stage are indicated by filled circles. The High Birth Rank subjects made on average 210.3 Errors to reach criterion as compared to 161.0 Errors for the Low-Rank subjects. However, this difference did not reach significance [F(1,18) =0.555, p=0.466]. *By Cog Rank*: Several animals changed ranks in adulthood and when using Cog Rank (see Figure 2B), 3 Low Cog rank animals failed to reach criterion on SD1 (filled circles). The High Cog Rank Group averaged 165.0 Errors and Low Cog Rank group averaged 189.1 Errors, a group difference that did not reach significance(F(1,18) = 0.35, p>0.05]. Thus, irrespective of their ranks at birth or at cognitive testing, performance on the simple discrimination did not differ between the High- and Low-ranking groups.

#### 4.2.2 Reversals (SR1-SR3)

Three subjects quit testing during this phase, two during SR1 (i.e., lost motivation/stopped working) and one during SR2 (after giving birth), therefore their data were excluded from analyses of this phase (data from these subjects in the SD stage are indicated by open squares in Fig. 2A, B. *By Birth Rank:* As shown in Figure 2A, from the 14 subjects (5 High; 9 Low), two that quit testing were Low-Ranking and one High-ranking (see open squares, Fig. 2A, SD tests). The remaining subjects completed the 3 Reversals. In general, most subjects improved across the three Reversals, regardless of Rank. The RM-ANOVA revealed no Main effect of Birth Rank [F(2,24)=0.44, p>0.05], or of Reversal [F(2,24)=1.86, p>0.05], nor was there a Birth Rank x Reversal interaction [F(2,24)=0.118, p>0.05]. *By Cog Rank:* As shown in Figure 2B, when the data from the 14 tested subjects were organized by Cog Rank (7 High; 7 Low), the Groups showed improved performance across the reversals [F(1,12)=1.95, p>0.05], but no group differences [F(1,12)=1.15, p>0.05] and no Cog-Rank x Reversal interaction [F(2,24) = 0.846, p>0.05]. *Perseverative Errors:* We compared the tendency to persist in choosing the incorrect stimulus (perseverations) during the three Reversals (data not shown).

Despite large individual differences in perseverations, on average, for either Birth or Cog Rank, subjects committed fewer perseverative errors with each reversal. RM-ANOVAs for Perseverations by Rank x Reversal did not yield any effects of Rank or Reversal [p’s>0.05]. Thus, the data for three reversals indicated similar performance for both the High- and Low-ranking subjects when analyzed either by their Birth of Cog-Ranks.

#### 4.2.3 ID/ED Shifts

The same 14 subjects that completed the Reversals were tested on the 3 ID/ED Shift stages (CD, ID, ED) and all subjects passed these 3 tests, except Sub-7 that could not attempt Stage 7 due to illness (its missing score for EDS was replaced with the mean of the Rank group for that test). Performance across the ID/ED Shifts are presented in Figure 2A & B, and data from the missing subject are indicated in the diamond markers (open for CD and IDS, filled gray for EDS). Considering individual scores, some subjects performed well from the beginning of this phase, others showed improvement across the three Shifts, and some showed no improvement. There were group differences depending upon whether Rank was determined by Birth or by Cog, as revealed by RM-ANOVAs across the Shift levels. *By Birth Rank*: As shown in Figure 2A using the 5 High and 9 Low animals, Low-ranking subjects committed fewer errors on average in the CD stage and showed about the same level of performance across the remaining two Shifts. By contrast, High-ranking subjects made more errors on the CD stage but then showed improvement across the ID and ED Shifts. RM-ANOVA confirms this description revealing no Main effect of Birth Rank [F(1,12)=0.004, p>0.05], a Main effect of Shifts [F*_Huynh-Feldt_* (1.44, 17.2)=6.32, p=0.014], and a Shift x Rank interaction [F*_Huynh-Feldt_* (1.44, 17.2)=5.45, p=0.022]. Pairwise comparisons (Bonferroni adjusted) of the effect of Shifts revealed that overall improvement across Shifts was reliable between the first (CD) and last (EDS) stages [p=0.036]. Further exploration of the interaction revealed that the performance improvement was largely limited to the High-Ranking group, in which there was reliable improvement from CD to IDS [p=0.034] and CD to EDS [p=0.023]. However, at each shift separately, the group difference did not reach significance. *By Cog Rank*: As shown in Figure 2B, when using the 7 High and 7 Low animals, both groups improved performance with each stage and showed similar Errors to Criterion. This description was confirmed with RM-ANOVA that yielded no Main Effect of Cog Rank [F(1,12)=0.419, p>0.05], or of Shift [F*_Huynh-Feldt_*(1.33,16.0)= 2.76, p>0.05], and no reliable Cog Rank x Shift interactions [F*_Huynh-Feldt_* (1.33, 16.0)= 0.08, p>0.05]. Therefore, the only difference we found when comparing shifts performance according to Birth or Cog ranks is that when using Birth rank, High-ranking animals show an improvement in performance from CD to ED and ID, but the Low-ranking animals did not.

### 4.3 Correlations between Rank and Personality

To assess whether Rank (Birth or Cog) correlated with personality type in either setting, we compared both individual traits and composite scores (as described above) with percentile ranks. Spearman Rank correlations on both Birth and Cog rank revealed associations only with Ob-trait scores. Both rank types correlated positively with Aggressive_Ob,_ suggesting that higher-ranking animals tended to be more aggressive in the social setting [rho = 0.438, p=.029 and rho = 0.621, p= 0.001, for Birth and Cog respectively]. Both Birth and Cog also correlated negatively with Anxious_Ob_ reflecting the tendency of lower-ranking subjects to display more anxiety in the social group setting {rho = −0.404, p=0.045 and rho=-0.517, p=0.008 for Birth and Cog respectively]. Birth rank further correlated positively with Curious_Ob_ [rho=0.423, p=0.035], reflecting higher levels of Curiosity in high-ranking subjects in the group setting. By contrast, Cog rank correlated negatively with Fearful_Ob_ [rho=-0.567, p=0.003], Timid_Ob_ [rho=-0.416, p=0.039], Gentle_Ob_ [rho=-0.455, p=0.022], and lastly with the composite Timid_Ob_ score [ rho=-0.442, p=0.027]; all reflecting the tendency of lower-ranking animals to be more Timid, Fearful and Anxious in the group setting.

### 4.4 Correlations between Performance, Birth vs Cog Rank, Personality

To determine whether Ranks and/or Personality traits correlated with performance in different phases of testing, we compared the traits listed in Supplementary Figure 3 across the Habituation/Pretraining phase and across the errors committed during the three testing phases of the ID/ED task as well as Birth and Cog Rank. Despite high correlations between the two Rank measures, and between some of the traits, not all correlated measures predicted performance equally.

#### 4.4.1 Habituation

For this phase, we used the Total Sessions to complete Habituation, and the Total Trials to pass Priming, for those subjects that passed this phase (to avoid artificial low scores for subjects that quit, rather than passed). As shown in Figure 3A, the number of sessions required to complete Habituation was reliably and negatively correlated with Cog Rank (but not Birth Rank), reflecting that Low Cog ranking animals took more sessions than High Cog ranking animals to habituate (rho = - 0.472, p = 0.023). With respect to personality traits (Figure 3B), there was a reliable, positive correlation between the Timid_Cog_ composite trait and Total Habituation Sessions, such that animals that scored higher on the Timid_Cog_ traits required more sessions to complete the Habituation phase (rho = 0.526, p=0.01). Interestingly, despite a high correlation of “Timid” (Cog- and Ob-traits, see Supplementary Figure 3), the Timid_Ob_ composite score did not predict Habituation performance. Lastly, trials to pass Priming did not show any relationship with Birth or Cog Rank, or Personality Traits.

**Figure 3:**
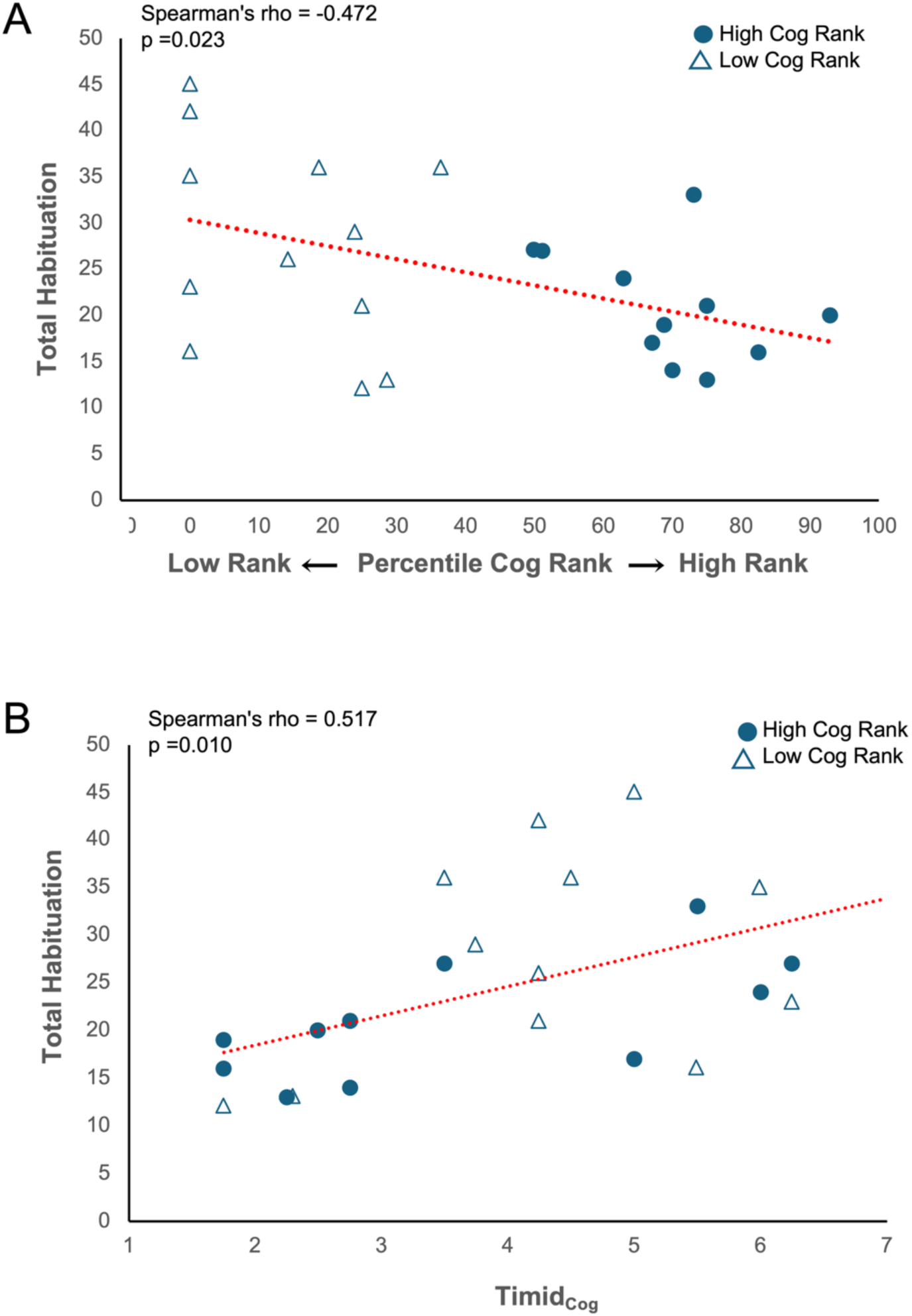
Habituation phase correlations. A: Total sessions to complete Habituation as a function of Cog Rank. B: Total sessions to complete Habituation as a function of Timid_Cog_ composite score (Likert scale 1-7). Blue circles indicate High Cog-ranking subjects, open triangles indicate Low Cog-ranking subjects

#### 4.4.2 ID/ED performance

For the three ID/ED phases, correlations were made on each stage separately. Errors committed in the SD1 stage had a moderate negative, but non-significant relationship with Curious_Ob_ [Spearman’s rho = −0.399, p=0.082]. For the *Reversals*, as shown in Figures 4A and B, there were no significant correlations between rank/personality and Reversal 1, whereas scores on Reversal 2 had a strong negative relationship with Timid_Cog_ [rho = −0.536, p=0.048], as well as with Timid_Ob_ that did not reach significance, but has a strong effect size [rho = −0.524, p=0.054], suggesting that very “Timid” subjects in the testing and social settings made fewer errors on this stage. Similarly, Errors on Reversal 3 (Fig. 4C) also showed a moderate, negative relationship with Timid_Cog_ (but not with Timid_Ob_) that did not reach significance [rho= −0.505, p=0.065]. For the *Shifts,* errors committed during performance of the ID (intradimensional) Shift were moderately-negative, but not reliably, correlated with Birth Rank [Spearman’s rho= −0.512, p=0.061], but not Cog Rank. Personality traits did not show any reliable relationship with Errors during the individual Shifts, however the Total Shift Errors showed a moderately-negative, but non-significant, correlation with Curious_Ob_ [rho=-0.503, p=0.067]. Each of those had moderately strong effect sizes.

**Figure 4:**
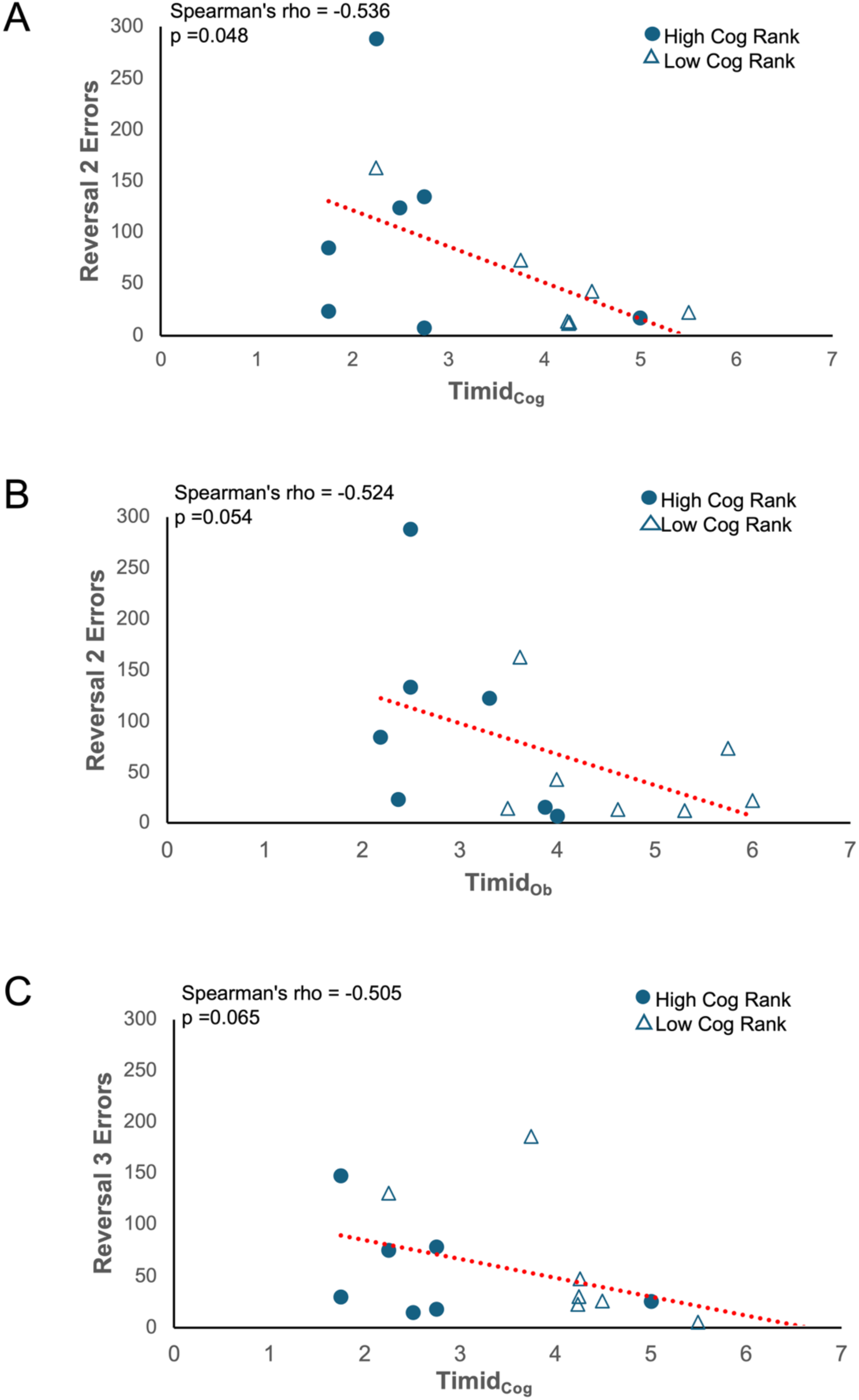
Reversal stage correlations: A) Reversal 2 correlations with Timid_Cog_, B) Reversal 2 correlations with Timid_Ob_, C) Reversal 3 Errors correlations with Timid_Cog_. Other conventions as in Figure 3.

## 5. DISCUSSION

In this study, we reported for the first time the cognitive performance of adult female rhesus macaques that have received chronic social subordination stress since birth. As they reached 7-8 years, subjects with high chronic social subordination stress and those with low social subordination stress were given the ID/ED task, a cognitive task that is relevant for age-related cognitive decline, particularly in executive function, and cognitive flexibility (Amelchenko et al., 2023; Bonté et al., 2011). We considered both Birth Rank, which is an indicator of the effects of early social subordination on the maturation of biological and neural measures but also Cog (current) Rank as some subjects have gained or lost social status due to changes in natal group sizes or family composition and which is a marker of cumulative effects of social stress at the time of testing.

Contrary to our expectation, we identify no significant changes in performance in the Low-ranking animals as compared to the high-ranking animals when using either the Birth Rank or Cog Rank, although the higher number of errors of some of the Low-ranking animals in the shifts portion of the task may already reflect the physiological and neural stress-related neural alterations previously reported in the Low-ranking animals (Kovacs-Balint et al., 2025; Kovacs-Balint et al., 2024a; Kovacs-Balint et al., 2024b). Interestingly, the impact of Birth Rank (and therefore High/Low Early Life Social Subordination Stress) was a better predictor of long-term consequences of social stress on task performance, whereas Cog (current) rank was somewhat more predictive of current temperament/personality, and how subjects interacted with the testing environment. These data will be discussed in greater detail below.

### 5.1 Effect of Rank on Habituation/Priming

These subjects are part of the ENPRC breeding colony and are therefore unaccustomed to removal from the social groups for anything other than clinical stays, or sample/neuroimaging collections. For higher ranking animals in particular, such removals are purposely short-term to maintain social stability of the group. As such, training them for cognitive testing proved to be a challenge. Rank played a role when it came to acclimating to the testing environment as a number of high-ranking animals were unsurprisingly less cooperative when out of their groups. For some of those, food rewards did not always provide sufficient enticement to cooperate with testing (as opposed to being in their social group). By contrast, access to high value treats in the absence of social competition was more of an inducement to many lower-ranking subjects. Thus, we had several high-ranking subjects that refused to engage in testing, as well as one high and one low-ranking subject that were fearful of the testing environment or touching the testing objects to pass priming.

Interestingly, we observed a negative relationship between Cog Rank and trials required to complete the pretraining phases, such that Higher-ranking animals required fewer sessions to complete the pretraining than lower ranking animals (Figure 3A). No significant relationships were seen when data were analyzed by Birth rank, indicating that the observed effects may represent a temperamental state of the animals at the time of testing and not of an early effect of chronic social subordination stress during the developmental period (Lupien et al., 2009).

### 5.2 Effects of Rank on ID/ED Testing

Social subordination stress starting early in life for Low-ranking females in this study was predicted to result in accelerated cognitive aging and cognitive decline, but that at 8 years of age, those effects may be mild and not significant or not yet evident. However, considering the physiological and neural changes we have already documented for these female monkeys between 8-10 years of age (Kovacs-Balint et al., 2025; Kovacs-Balint et al., 2024a; Kovacs-Balint et al., 2024b), one might expect earlier behavioral and cognitive signs of those changes. Yet, we found no group differences of Birth or Cog Rank in performance on any of the stages of the ID/ED task. Nevertheless, inspection of the averaged scores of the groups (Figure 2A) indicates that Low Ranked animals did not differ from High Ranked in terms of errors to criterion for the SD, or Reversal stages. They did make slightly more errors in the IDS and EDS stages and also showed little improvement across the three Shifts. By contrast, High Ranked females showed the greatest improvement in the Shift stages of the ID/ED task. Yet, this slight group differences depended on their Birth Rank, not their Cog Rank (Fig. 2B). This suggests that the relevant impact of rank on task performance resulted from the effects of social stress that began during the early developmental period, rather than to the effects of current chronic stress at the time of testing (Lupien et al., 2009).

### 5.3 Personality traits correlations with Rank and Performance

The role played by personality was also reflected by Rank. We found similar correlations whether by Birth or Cog rank with Aggressive and Anxious in the Ob (group social observations) setting, with higher rank predicting more aggression and less anxiety. Birth rank also predicted Curious in the Ob setting. However, Cog Rank, but not Birth Rank, was predictive of Fearful, Timid and Gentle ratings for Low ranking subjects in the Ob setting. With respect to the impact of personality traits on performance, Total Habituation had a negative relationship with Cog Rank (higher rank, faster learning) and a positive relationship with Timid_Cog_ (higher Timid rating, slower learning). As for performance on the ID/ED task, errors on Reversals 2 were negatively correlated with Timid_Cog_ (higher Timid rating = fewer errors). Nonsignificant correlations were also found for Timid_Ob_ and Timid_Cog_ with Reversal 2 and Reversal 3 Errors, respectively. Interestingly, no personality trait was correlated with Errors on the Shifts. In general, personality traits played a greater role in the pretraining phases of cognitive testing, that is, whether a subject was too anxious or more curious could impact their willingness to acclimate with the testing environment and to engage in testing. These traits did not appear to impact cognitive ability, however the relationship of the Timid composite scores, which includes Anxious, Fearful, Cautious and Timid traits did relate to performance on Reversal 2, presumably reflecting some increased level of frustration during the reversal stages.

### 5.4 Cognitive Performance Correlations with Neural Changes

The impact of chronic social subordination stress in our subjects has also changed a number of physiological measures not reported here. Of particular interest for cognitive performance is the impact of high social stress to brain development and aging. In recent neuroimaging assessments of these animals (at the same age as cognitive testing), we found structural and functional connectivity changes in the PFC-HIPP-AMY networks in Low Birth Rank subjects that also correlated with some aspects of cognitive performance (Kovacs-Balint et al., 2025; Kovacs-Balint et al., 2024a; Kovacs-Balint et al., 2024b). Our preliminary data show that despite limited effects of rank on cognition, neural differences in SUBs correlated with errors in specific stages of the ID/ED task. For example, larger HIPP volume in SUBs predicted more IDS errors (rho=0.525, p=0.05), presumably due to a dysfunctional medioventral PFC to which the HIPP projects (Aggleton et al., 2015) and when damaged affects IDS performance (Dias et al., 1996). In addition, larger AMY volume predicted more errors in Reversal 2 (rho=0.556, p=0.039), presumably due to a dysfunctional ventrolateral PFC, including the ORB cortex, to which the AMY projects (Aggleton et al., 2015; Ghashghaei et al 2007) and when damaged impairs Reversal performance (Dias et al., 1996). Further analyses with our entire sample size will likely strengthen these associations. These findings indicate that structural brain changes precede the emergence of cognitive changes.

### 5.5 General Discussion

One of the most fundamental and urgent goals of research into the cognitive neuroscience of aging remain to better understand why some individuals decline faster than others during healthy aging. The present study was conducted to determine the impact of early life and chronic social subordination stress to accelerated cognitive aging in macaques. Although one would not predict age related performance differences at the adult age of 8 years, the measurable changes in other physiological and neural measures in these animals suggest that there have either been differences beginning early in life in programming brain development or resulting from cumulative exposure to subordination stress. However, that we found minimal impact on cognitive ability so far is interesting. We would expect increasing differences as these animals reach middle age, with the SUB group showing declining performance at an earlier age than DOMs. Another interesting finding that came out of this investigation is that performance of the animals in almost each stage of the task showed important individual differences similar to those described in the earlier reports (Baxter et al., 2023; Lizarraga et al., 2020; Rapp and Amaral, 1992; Sadoun et al., 2019; Vanderlip et al., 2024; Vanderlip et al., 2023; Varma et al., 2024; Zeamer et al., 2011). This individual variability in the scores could be seen in both groups, whether using Birth Rank or Cog Rank, that is the DOM group showed as much individual variability between the animals’ scores as the SUB group that had received chronic social stress since birth. Thus, the chronic social stress may not be the most critical factor that may explain the variability in the scores of the aged animals of the previous studies in which the social status of the animals was not available. However, inter-individual variability in cognitive aging is striking even in normal aging humans (Habib et al., 2007) and is likely to reflect a complex interaction between genetic and environmental factors (Cabeza et al., 2018). The impact of “Rank”, in fact may differ across individuals and experiences. For example, Abbott and colleagues (Abbott et al., 2003) have suggested that not all NHP individuals of low social rank are always stressed. For example, those that have fewer sources of social support, and greater frequency of stressors will show higher levels of hypercortisolism than those that receive more buffer from their social group. That would suggest we might still observe differences in cognitive decline even within the SUB group depending on the balance of stressors and social support they have experienced in their lifetime.

### 5.6 Limitations

One of the advantages of nonhuman primates such as Rhesus monkeys is their long life spans that can show gradual emergence of aging-related cognitive impairments (e.g. in cognitive flexibility, executive functions, memory) with similar time courses as humans (e.g., emerging in middle age, and with parallel age-related neural changes (e.g. myelin loss, cortical thinning, altered connectivity, neuropathology) in prefrontal cortex (PFC), amygdala (AMY), and hippocampus (HIPP). Thus, the use of NHPs is clinically relevant given that the impacts of early life and chronic stress in developing humans has long been of interest. However, this study also has several limitations to generalizing the results. For example, we had sample size reductions due to difficulty testing some of the animals, or attrition due to birth and care of infants during the months of cognitive testing, (although that is a translational experience for human mothers). However, planned further longitudinal testing as these animals approach middle- and pre-geriatric ages, will determine in the same population the trajectory of cognitive decline. We will include additional cognitive tasks that may tap into other aspects of cognitive function. The impact of lifetime Rank changes is more difficult to interpret in this instance as those changes resulted from group divisions that gave opportunity for Low-ranking animals to improve their social standing, or to cause High-ranking individuals to lose standing when former lower-ranked members change groups. Those events generally occurred between puberty and adulthood, as these subjects were born and reared in large (∼180 individuals) multifamily social groups. Thus, the effects of rank-changes/Cog Rank due to group splintering, which occurred largely after menarche, may have had a different impact than the effect of Birth Rank alone. That is, the role of early social subordination may have “programming” effects on brain development as opposed to the cumulative consequences of chronic social stress that occurred across adulthood. Our results are in line with this hypothesis as we observed in our testing that Birth Rank was the better predictor of cognitive performance differences, limited though they were. This finding agrees with the impact of early stress on the developing brain. For example, hippocampal development is especially vulnerable to glucocorticoids surges in the first two years of life in humans (Lupien et al., 1998) (∼first 6 months in macaques). Cog Rank was a better predictor of personality traits assessed in adulthood at the age of cognitive testing

Lastly, is the limitation of using only females in a translational model of social stress and aging. Despite differences in longevity, risk for age related dementias and responses to stress, much of the early animal aging studies were conducted in male subjects. For example, in humans, the impacts of early life stress may differ between the sexes, with human males being more vulnerable to early life stress/adversity in childhood, but in human females it becomes more pronounced with time (Logue and Nemeroff, 2025). In macaques, in addition to the model of social subordination largely impacting females, they were chosen because in humans, women live longer on average than men, and as such are at higher risk for normal age-related changes in cognition as well as dementias, and rhesus macaques naturally live in predominantly female, multi-family social groups. However, given that differences in responses to early life stress are evident before puberty in humans, and/or may be independent of hormonal surges (Birnie and Baram, 2025; Lupien et al., 2018; Lupien et al., 2009), and similarly, cognitive aging impairments emerge earlier in females than males in marmosets (Rothwell et al., 2022), thus the present results may not transfer well to models of male aging in monkeys or humans.

## 6. Conclusions

The present study provides further evidence that chronic social subordination stress beginning early in life not only alters physiological stress-related measures and developmental trajectories of brain structures that are maturing from infancy through early adulthood (Godfrey et al., 2018; Kovacs-Balint et al., 2025; Kovacs-Balint et al., 2024a; Kovacs-Balint et al., 2024b; Kyle et al., 2019) but can also be expected to play a role in the cumulative effects of chronic stress throughout life. As they have reach full adulthood, females with high levels of social subordination stress showed only minor changes in attentional flexibility, despite measurable differences in brain structures associated with cognitive function. Planned longitudinal testing as they reach pre-geriatic stage and older may reveal differences in the emergence of cognitive aging as an effect of social subordination stress.

## ACKNOWLEDGEMENTS

This study was conducted with invaluable help animal care, colony management and veterinary staff at the Emory National Primate Research Center (ENPRC) Field Station. This project was funded by NIH grants AG070704, HD077623, the NIH’s Office of the Director, Office of Research Infrastructure Programs P51OD011132 (ENPRC Base Grant), and the EPC Fund for Excellence. The ENPRC is fully accredited by AAALAC, International.

