## Supplementary Figures for "The effects of chronic social stress on cognitive flexibility in adult female macaques"

| | | Fearful$_{Cog}$ | Cautious$_{Cog}$ | Timid$_{Cog}$ | Anxious$_{Cog}$ | Gentle$_{Cog}$ | Cool$_{Cog}$ | Curious$_{Cog}$ | Aggressive$_{Cog}$ | Irritable$_{Cog}$ | Defiant$_{Cog}$ | Excitable$_{Cog}$ | Erratic$_{Cog}$ |
|---|---|---|---|---|---|---|---|---|---|---|---|---|---|
| Fearful$_{Cog}$ | | | | | | | | | | | | | |
| Cautious$_{Cog}$ | *rho* | 0.876 | | | | | | | | | | | |
| | *p* | 0.000 | | | | | | | | | | | |
| | *N* | 25 | | | | | | | | | | | |
| Timid$_{Cog}$ | *rho* | 0.910 | 0.866 | | | | | | | | | | |
| | *p* | 0.000 | 0.000 | | | | | | | | | | |
| | *N* | 25 | 25 | | | | | | | | | | |
| Anxious$_{Cog}$ | *rho* | 0.691 | 0.638 | 0.752 | | | | | | | | | |
| | *p* | 0.000 | 0.001 | 0.000 | | | | | | | | | |
| | *N* | 25 | 25 | 25 | | | | | | | | | |
| Gentle$_{Cog}$ | *rho* | 0.464 | 0.439 | 0.409 | -0.016 | | | | | | | | |
| | *p* | 0.034 | 0.047 | | | | | | | | | | |
| | *N* | 21 | 21 | | | | | | | | | | |
| Cool$_{Cog}$ | *rho* | -0.168 | -0.108 | -0.159 | -0.460 | 0.840 | | | | | | | |
| | *p* | | | | 0.024 | 0.000 | | | | | | | |
| | *N* | | | | 24 | 21 | | | | | | | |
| Curious$_{Cog}$ | *rho* | -0.671 | -0.572 | -0.551 | -0.573 | 0.104 | 0.462 | | | | | | |
| | *p* | 0.000 | 0.004 | 0.005 | 0.003 | | 0.023 | | | | | | |
| | *N* | 24 | 24 | 24 | 24 | | 24 | | | | | | |
| Aggressive$_{Cog}$ | *rho* | -0.498 | -0.575 | -0.494 | -0.114 | -0.781 | -0.359 | 0.006 | | | | | |
| | *p* | 0.011 | 0.003 | 0.012 | | 0.000 | | | | | | | |
| | *N* | 25 | 25 | 25 | | 21 | | | | | | | |
| Irritable$_{Cog}$ | *rho* | -0.430 | -0.468 | -0.332 | -0.075 | -0.476 | -0.095 | 0.038 | 0.725 | | | | |
| | *p* | 0.036 | 0.021 | | | 0.029 | | | 0.000 | | | | |
| | *N* | 24 | 24 | | | 21 | | | 24 | | | | |
| Defiant$_{Cog}$ | *rho* | -0.240 | -0.306 | -0.304 | 0.036 | -0.575 | -0.377 | -0.113 | 0.640 | 0.436 | | | |
| | *p* | | | | | 0.006 | | | 0.001 | 0.033 | | | |
| | *N* | | | | | 21 | | | 24 | 24 | | | |
| Excitable$_{Cog}$ | *rho* | -0.326 | -0.301 | -0.342 | 0.055 | -0.723 | -0.529 | -0.100 | 0.708 | 0.607 | 0.717 | | |
| | *p* | | | | | 0.000 | 0.009 | | 0.000 | 0.002 | 0.000 | | |
| | *N* | | | | | 21 | 23 | | 23 | 23 | 23 | | |
| Erratic$_{Cog}$ | *rho* | -0.129 | -0.175 | -0.082 | 0.147 | -0.234 | 0.064 | -0.141 | 0.484 | 0.757 | 0.254 | 0.448 | |
| | *p* | | | | | | | | 0.017 | 0.000 | | 0.032 | |
| | *N* | | | | | | | | 24 | 24 | | 23 | |

**Supplementary Figure 1**: Correlation matrices for personality traits: Traits measured at Cognitive Testing. Spearman Rank Correlations between traits rated in the Cog post-testing condition. Red shading indicates significant positive correlations, blue shading indicates significant negative correlations. Clusters of correlated traits suggested across the Fearful, Cautious, Timid and Anxious traits ("Timid"); across the Aggressive, Irritable, Defiant, Excitable and Erratic traits ("Bold"); and across the Gentle, Cool and Curious traits ("Calm).

| | | Fearful$_{Ob}$ | Cautious$_{Ob}$ | Timid$_{Ob}$ | Anxious$_{Ob}$ | Gentle$_{Ob}$ | Cool$_{Ob}$ | Curious$_{Ob}$ | Aggressive$_{Ob}$ | Irritable$_{Ob}$ | Defiant$_{Ob}$ | Excitable$_{Ob}$ | Erratic$_{Ob}$ |
|---|---|---|---|---|---|---|---|---|---|---|---|---|---|
| Fearful$_{Ob}$ | | | | | | | | | | | | | |
| Cautious$_{Ob}$ | *rho* | 0.761 | | | | | | | | | | | |
| | *p* | 0.001 | | | | | | | | | | | |
| | *N* | 25 | | | | | | | | | | | |
| Timid$_{Ob}$ | *rho* | 0.808 | 0.830 | | | | | | | | | | |
| | *p* | 0.001 | 0.001 | | | | | | | | | | |
| | *N* | 25 | 25 | | | | | | | | | | |
| Anxious$_{Ob}$ | *rho* | 0.783 | 0.645 | 0.693 | | | | | | | | | |
| | *p* | 0.001 | 0.001 | 0.001 | | | | | | | | | |
| | *N* | 25 | 25 | 25 | | | | | | | | | |
| Gentle$_{Ob}$ | *rho* | 0.355 | 0.303 | 0.227 | 0.115 | | | | | | | | |
| | *p* | | | | | | | | | | | | |
| | *N* | | | | | | | | | | | | |
| Cool$_{Ob}$ | *rho* | -0.307 | -0.213 | -0.244 | -0.507 | 0.107 | | | | | | | |
| | *p* | | | | 0.01 | | | | | | | | |
| | *N* | | | | 25 | | | | | | | | |
| Curious$_{Ob}$ | *rho* | -0.309 | -0.104 | -0.378 | -0.239 | 0.102 | -0.046 | | | | | | |
| | *p* | | | | | | | | | | | | |
| | *N* | | | | | | | | | | | | |
| Aggressive$_{Ob}$ | *rho* | -0.624 | -0.384 | -0.627 | -0.486 | -0.404 | 0.125 | 0.332 | | | | | |
| | *p* | 0.001 | | 0.001 | 0.014 | 0.05 | | | | | | | |
| | *N* | 25 | | 25 | 25 | 25 | | | | | | | |
| Irritable$_{Ob}$ | *rho* | -0.179 | -0.243 | -0.144 | -0.012 | -0.536 | 0.049 | -0.048 | 0.450 | | | | |
| | *p* | | | | | 0.006 | | | 0.024 | | | | |
| | *N* | | | | | 25 | | | 25 | | | | |
| Defiant$_{Ob}$ | *rho* | -0.283 | -0.347 | -0.330 | -0.149 | -0.303 | 0.025 | -0.075 | 0.471 | 0.390 | | | |
| | *p* | | | | | | | | 0.017 | | | | |
| | *N* | | | | | | | | 25 | | | | |
| ExcitableOb | *rho* | 0.071 | 0.050 | -0.036 | 0.395 | -0.106 | -0.692 | 0.321 | 0.195 | 0.144 | 0.044 | | |
| | *p* | | | | | | 0.001 | | | | | | |
| | *N* | | | | | | 25 | | | | | | |
| ErraticOb | *rho* | -0.064 | 0.023 | -0.045 | 0.262 | -0.360 | -0.284 | 0.273 | 0.484 | 0.617 | 0.310 | 0.551 | |
| | *p* | | | | | | | | 0.014 | 0.001 | | 0.004 | |
| | *N* | | | | | | | | 025 | 25 | | 25 | |

**Supplementary Figure 2**: Correlation matrices for personality traits: Traits measured in Social Group. Spearman Rank Correlations between traits rated in the Ob social condition. Red shading indicates significant positive correlations, blue shading indicates significant negative correlations. For this condition, the same "Timid" cluster is highly correlated, and a less highly correlated cluster can be made for "Bold", excluding the Excitable trait. All other traits will be used in the model as individual predictors.

| | | **Timid_Cog** | **Calm_Cog** | **Bold_cog** |
|---|---|---|---|---|
| **Timid_Ob** | *rho* | 0.52 | -0.12 | -0.16 |
| | *p* | 0.008 | | |
| | *N* | 25 | | |
| **Bold_Ob** | *rho* | -0.28 | -0.36 | 0.15 |
| | *p* | | | |
| | *N* | | | |
| Gentle_ob | *rho* | 0.26 | 0.42 | -0.39 |
| | *p* | | 0.039 | |
| | *N* | | 24 | |
| Cool_ob | *rho* | 0.20 | -0.01 | -0.18 |
| | *p* | | | |
| | *N* | | | |
| Curious_ob | *rho* | -0.33 | 0.35 | -0.26 |
| | *p* | | | |
| | *N* | | | |
| Excitable_ob | *rho* | -0.31 | 0.14 | 0.05 |
| | *p* | | | |
| | *N* | | | |

**Supplementary Figure 3**: Correlation matrices for personality traits: Correlations for Cognitive Testing vs. Social Group Personality Traits. Spearman Rank Correlations across Cog and Ob ratings utilizing the composite scores as described abpve as well as the individual Ob predictors. Composite scores are further indicated in bold text. The Timid traits correlated between the Cog and Ob settings, as did Gentle ratings in the Social Ob setting with the Calm composite score in the testing setting.
